# Rare sex punctuates strict asexual reproduction in the clonal raider ant, *Ooceraea biroi*

**DOI:** 10.64898/2026.07.01.735869

**Authors:** Kip D. Lacy, Nicolas Châline, Daniel J.C. Kronauer

**Affiliations:** Laboratory of Social Evolution and Behavior, The Rockefeller University, New York, NY, USA; Institute of Psychology, University of São Paulo, São Paulo, Brazil; Howard Hughes Medical Institute, New York, NY, USA

## Abstract

While asexual species can often outcompete their sexual counterparts over ecological timescales, their long-term evolutionary success is hindered by a diminished ability to purge deleterious mutations and to adapt to changing environments. However, some asexual species persist for millions of years, and a major question in evolutionary biology is how they do so. One solution is to occasionally reproduce sexually, as has been shown in a handful of primarily asexual species. Here, we investigate the possibility of rare sex in the clonal raider ant, *Ooceraea biroi*. We report the whole-genome sequence of a previously uncharacterized clonal line and, using population genetic and phylogenetic analyses, show that it originated through sexual reproduction between two extensively studied clonal lines. The mitochondrial genome of this clonal line differs from that of the maternal clonal line at only a single nucleotide, suggesting that the sexual reproduction event occurred within the past few hundred years. These results demonstrate that sex occurs sporadically in the clonal raider ant, allowing it to generate new genetic combinations and potentially to overcome some of the costs of asexuality.

## Introduction

A longstanding puzzle in evolutionary biology is how asexual reproduction can persist over long periods of time. Meiosis and sex likely evolved before the last common ancestor of eukaryotes and are maintained in the vast majority of animal species [1]. However, asexual species evolve sporadically [2,3]. Asexual reproduction can be more efficient than sexual reproduction in the short term, leading to ecological success [4,5], which is reflected in the invasive potential of many asexual species [6–8]. Despite these ecological advantages, asexual species are thought to be evolutionarily short-lived because asexuality leads to reduced genetic diversity and reduced ability to purge deleterious mutations, resulting in long-term costs and, ultimately, extinction [2,9–11]. Sex, on the other hand, generates new combinations of alleles that are exposed to natural selection, allowing deleterious alleles to be selectively eliminated [9,12] and facilitating adaptive evolution [13,14], for example, in response to parasites and other ecological threats [15,16].

However, asexual reproduction can persist, sometimes over millions of years [2,17]. In principle, rare sexual reproduction in otherwise strictly asexual lineages could sporadically introduce genetic diversity and ameliorate some of the costs of asexuality [18,19]. Recent studies using genetic data have lent credence to this idea, with rare sex detected in species previously thought to reproduce exclusively asexually, including brine shrimp [20], bdelloid rotifers [21–23], and *Timema* stick insects [24].

Here, we use whole-genome sequencing to study the possibility of rare sex in the clonal raider ant, *Ooceraea biroi*. This species reproduces asexually [25], employing a form of thelytokous (female-producing) parthenogenesis (development from an ovum without fertilization by sperm) called “automixis with central fusion,” in which the two central meiotic products fuse after meiosis II [26,27]. This mode of reproduction is expected to lead to loss of heterozygosity [28,29]. However, *O. biroi* maintains heterozygosity with high fidelity due to non-Mendelian co-inheritance of recombined chromatids [30]. Unlike most ant species, in which queens mate and reproduce while workers perform other tasks and lay few, if any, eggs, *O. biroi* lacks queens [25]. Thus, colonies are composed entirely of workers that reproduce asexually and are genetically nearly identical [27,30–32]. Native populations of *O. biroi* have been found in Bangladesh, but the species has become introduced on tropical and subtropical islands around the world [26,33,34].

Ants, like all Hymenoptera, are haplodiploid, meaning that females are diploid and males are haploid. Haploid males occur rarely in laboratory colonies of *O. biroi* and likely result from rare failures of central fusion after meiosis [26]. In principle, these rare haploid males could enable *O. biroi* to occasionally reproduce sexually. However, we have not found any signs of sexual reproduction in over 17 years of keeping the species in the lab in large numbers.

Rare sex between otherwise purely clonal lines of *O. biroi* was previously hypothesized based on genotyping data at several microsatellite loci [26]. However, sexual reproduction has been impossible to infer definitively due to limited data. Here, we report the whole-genome sequence of an individual from clonal line T4, of which a single colony was collected in Taiwan in 2001, where it co-occurred alongside clonal lines A and B (**figure 1a**). Based on microsatellite data, T4 appeared to be intermediate between the extensively studied clonal lines A and B (Kronauer et al., 2012). By comparing this new genome to published genome sequences from other clonal lines, we find that clonal line T4 is most likely derived from a sexual reproduction event between a clonal line A female and a clonal line B male, demonstrating that rare sexual reproduction indeed occurs in *O. biroi*.

**figure 1.**
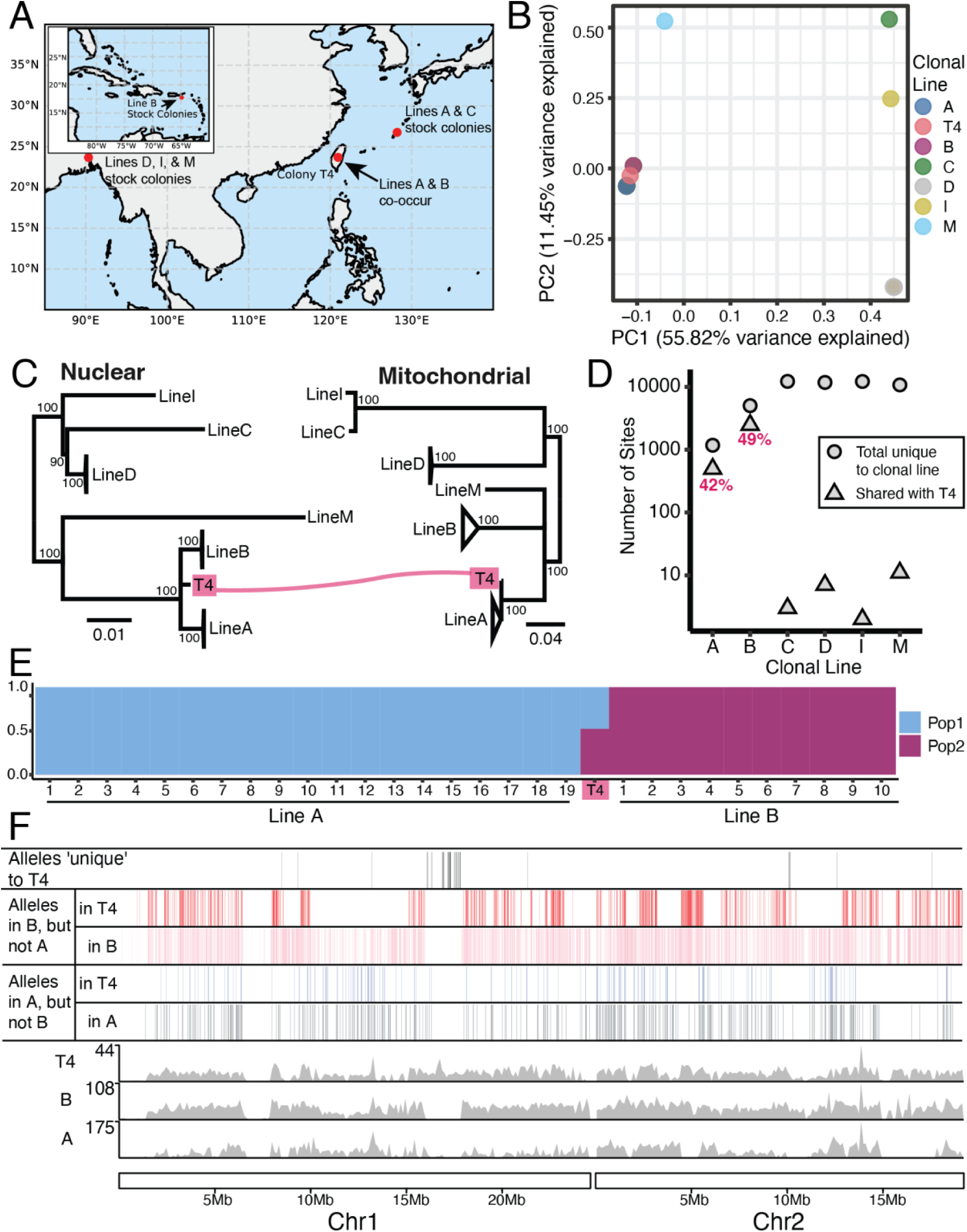
An *O. biroi* clonal line collected in Taiwan resulted from sex between two other clonal lines. (**a**) Partial maps of East Asia and the Caribbean depicting the collection locales of different colonies of *O. biroi*. Note that the sequenced individuals from clonal lines A and B came from colonies originally collected on Okinawa and St. Croix, respectively, rather than Taiwan. (**b**) Principal components analysis of SNP variation in *O. biroi*. (**c**) Midpoint-rooted maximum likelihood phylogenies based on SNPs from nuclear and mitochondrial genomes. Bootstrap supports from 1000 bootstrap replicates for major nodes are shown, and nodes with bootstrap support <90 were collapsed. Scale bars indicate substitutions per SNP. Phylogenies showing all individual samples are provided in **figure S1**. (**d**) The number of sites with alleles that, excluding T4, are unique to each clonal line (private alleles) and, among those sites, the number of sites at which the private allele is shared with T4. For clonal lines A and B, the proportion of sites shared with T4 is shown in red. (**e**) Stacked bar plot depicting the proportional assignment of each individual from clonal lines A and B, as well as the individual from colony T4, to two genetic clusters inferred from the admixture analysis (metadata for all DNA sequencing libraries included in this study are in **table S1**). (**f**) Karyoplot depicting, for the first two chromosomes in the *O. biroi* genome, sites with informative alleles in clonal lines A, B, and T4 as vertical tick marks. The numbers of heterozygous sites per 120kb window are shown in the gray histograms. Plots for all chromosomes are shown in **figure S2**.

## Methods

### Short-read whole-genome shotgun sequencing

We sampled a single adult worker from clonal line T4, disrupted the tissue using a Qiagen TissueLyser II, and extracted genomic DNA using Qiagen’s QIAmp DNA Micro Kit. We prepared a sequencing library using Illumina’s Nextera DNA Flex kit and sequenced on an Illumina NovaSeq 6000. Similar to other published *O. biroi* genomes of individual ants that we used for comparison [30,35,36], we targeted 40x sequencing coverage and achieved 42x coverage, ensuring sufficient coverage to detect heterozygosity accurately. We trimmed reads using Trimmomatic 0.36 [37], aligned them to the *O. biroi* reference genome (Obir_v5.4, GenBank assembly accession: GCA_003672135.1) using bwa mem [38], and subsequently sorted, deduplicated, and indexed using picard (http://broadinstitute.github.io/picard/). The metadata for all DNA sequencing libraries included in this study are in **table S1**.

We called variants using GATK HaplotypeCaller (version 4.2) [39] and filtered using GATK’s hard-filtering recommendations. We performed additional filtering to include only high-confidence variants in this analysis. We excluded falsely collapsed regions in the reference genome by filtering out sites found to be heterozygous in haploid males, variants with three or more alleles in a single diploid clonal line, and sites with read depth greater than twice the genome-wide mean. To exclude erroneous calls of heterozygosity or homozygosity, we filtered out sites with read depth of less than 15, with proportionate minor allelic depth of less than 0.25 in putatively heterozygous samples, or with nonzero minor allelic depth in putatively homozygous samples. These additional filtering steps are essential for obtaining a high-quality set of variants. In a previous study, we observed that forgoing any of these additional filtering steps led to many false positive gains of heterozygosity, losses of heterozygosity, and recombination events [30].

### Population genetic analyses

We used PLINK for principal components analysis [40] and ADMIXTURE v1.3 [41] to investigate population structure. To reconstruct maximum likelihood phylogenies, we converted VCFs to phylip format using vcf2phylip.py (https://github.com/joanam/scripts/blob/master/vcf2phylip.py) and used iqtree2 to reconstruct the phylogenies with the following arguments: “-st DNA -m TEST -bb 1000” [42–44].

### Data visualization

We used cartopy (https://cartopy.readthedocs.io) to draw maps, FigTree v1.4.4 (https://tree.bio.ed.ac.uk/software/figtree/) to draw phylogenies, and karyoploteR [45] to draw karyoplots. Other plots were drawn using ggplot2 [46] or Matplotlib [47]. Illustrations and final figures were made using Adobe Illustrator.

## Results

To investigate the possibility of sexual reproduction, we first conducted a principal components analysis using nuclear single nucleotide polymorphism (SNP) variation among all sequenced *O. biroi* individuals. These included several individuals from clonal line A collected on Okinawa, and of clonal line B collected on St. Croix (**figure 1a**; Kronauer et al. 2012). Unfortunately, samples of clonal lines A and B from Taiwan were not available for this study. The T4 individual clustered between clonal lines A and B along the first two principal components (**figure 1b**), indicating that it shares alleles with both clonal lines.

In sexual species, the nuclear genome is inherited from both parents, while the mitochondrial genome is only inherited from the mother. The pedigrees of mitochondrial and nuclear loci can therefore be discordant. However, in purely asexually reproducing lineages, where all genetic material is exclusively inherited from the mother, the evolutionary histories of the mitochondrial and nuclear genomes are entirely concordant. Therefore, discordance between nuclear and mitochondrial phylogenies can reveal sexual reproduction in otherwise asexual species [48]. Thus, we reconstructed maximum likelihood phylogenies separately using nuclear and mitochondrial SNP variation. The genome of the T4 individual occupies a separate branch in the nuclear phylogeny. However, it differs by only one SNP from the rest of the clonal line A polytomy in the mitochondrial phylogeny, whereas it differs by 271 SNPs from the clonal line B branch (**figure 1c**, **figure S1**). This further supports a sexual origin for clonal line T4. The fact that clonal lines A and T4 have nearly identical mitochondrial genomes implies that an individual from clonal line A was the mother of the individual that gave rise to clonal line T4, while an individual from clonal line B was its father.

To explore the nature of the relationship between clonal lines A, B, and T4 further, we excluded the T4 individual from the dataset and then identified sites with “private” alleles found exclusively in each clonal line. Then, for each clonal line, we counted the number of sites with otherwise private alleles that were also found in the T4 individual. Clonal lines A and B shared 42% and 49% of their otherwise private alleles with the T4 individual, respectively (**figure 1d**). These proportions are close to 50%, the pattern that would be expected if one of the sequenced individuals from each of those clonal lines was a parent of clonal line T4. The deviation from 50% likely results, in part, from genetic differences between our sequenced isolates of clonal lines A and B (which were collected from Okinawa and St. Croix, respectively) and the isolates of clonal lines A and B that produced T4 (which likely occurred in Taiwan).

The identity of clonal line T4 as a “daughter” line of clonal lines A and B was further supported by an admixture analysis. For this analysis, we included samples from clonal lines A, B, and T4, but unbiased analysis revealed that the optimal number of genetic clusters was two (**table S2**). Under this model, all individuals from clonal lines A and B were assigned entirely to their respective cluster, whereas the T4 individual was evenly assigned to the two clusters (47.6% to Pop1 and 52.4% to Pop2; **figure 1e**).

To examine the pattern of inheritance across the genome, we visualized the alleles that are private to clonal lines A and B along the reference genome and determined whether these alleles were also present in the T4 individual (**figure 1f, figure S2, figure S3**). There are two major takeaways from this analysis: (1) The T4 individual carries alleles that are otherwise private to either clonal line A or B on every chromosome, indicating that clonal line T4 inherited one version of each homologous chromosome from clonal line A and one version from clonal line B (**figure S3)**. (2) Within any genomic region inherited from clonal line A or B, the T4 individual inherited only a subset of the private alleles rather than all of them (**figure 1f, figure S2, figure S3**). This pattern is consistent with homologous recombination (usually without loss of heterozygosity; see [30]) over many generations in the ancestors of the mother and father of T4. Such recombination would have made the parental chromosomes transmitted to T4 shuffled versions of the ancestral haplotypes of each parental clonal line. Thus, when only one homologous chromosome from each parent was transmitted via sexual reproduction, T4 inherited only a subset of the private alleles.

In a smaller set of genomic regions, the T4 individual lacks alleles private to clonal line A or B, or it has alleles that have so far not been found in clonal lines A or B (alleles “private” to T4). Both patterns are consistent with the fact that our sequenced isolates of clonal lines A and B are not from Taiwan and with the described recombination dynamics in *O. biroi*. Although heterozygosity is usually maintained, crossovers can produce localized loss of heterozygosity at one or a few SNPs through gene conversion [30], and crossovers occasionally also lead to loss of heterozygosity across large segments of chromosomes [26,27,30,35,36]. The genomic regions with alleles “private” to T4 coincide with regions of homozygosity in our sequenced clonal line A or B individuals (**figure 1f, figure S2, figure S3**). Therefore, these alleles likely represent regions of the genome in which the parent of T4 (i.e., line A or B isolates from Taiwan from which we do not have sequenced genomes) retained heterozygosity that was lost in our sequenced isolates of clonal lines A and B (the isolates from Okinawa and St. Croix, respectively). Conversely, there are also genomic regions where the T4 individual possesses none of the private alleles from clonal line A and other regions where T4 possesses none of the private alleles from clonal line B (**figure 1f, figure S2, figure S3**). These likely represent losses of heterozygosity that occurred in the T4 lineage following the sexual reproduction event that produced the ancestor of the sequenced T4 individual.

The mitochondrial genome of T4 differs from that of the sequenced clonal line A individuals by a single SNP, allowing us to calibrate an estimate of how recently this sexual reproduction event occurred. The minimum estimate would be just prior to the collection of the colony in 2001 [26] because it is possible that this SNP difference already existed in the clonal line A mother of T4. However, it is also possible that the SNP difference arose via mutation after the sexual reproduction event, and we can estimate the upper bound by calculating the amount of time for a mitochondrial mutation to occur. To our knowledge, the mitochondrial genome mutation rate has not been directly measured for any hymenopteran. However, a mutation accumulation study in *Drosophila melanogaster* found a mitochondrial mutation rate of 6.3 * 10^-8^ per site per fly per generation (95% CI: (2.6 * 10^-8^, 13.4 * 10^-8^)) [49]. Assuming that *O. biroi* has a similar mitochondrial mutation rate to *Drosophila* and that there are six generations per year (as occurs under laboratory conditions), then one new mitochondrial mutation would be expected every 70-370 years, indicating that the sexual reproduction event that produced clonal line T4 likely occurred sometime in the past few hundred years.

## Discussion

Many animal lineages that were previously thought to reproduce entirely asexually show signs of rare sexual reproduction [20–24]. Here, we discovered a similar pattern in *O. biroi*. By leveraging a combination of phylogenetic and population genetic analyses of the nuclear and mitochondrial genomes of an individual from clonal line T4, which was originally collected in Taiwan, we found robust evidence that a clonal ancestor of that individual was produced by sexual recombination between a mother from clonal line A and a father from clonal line B (**figure 1**). This has likely occurred in the past few hundred years, revealing that although *O. biroi* reproduces almost entirely asexually, rare instances of sexual reproduction can occur.

The exact circumstances of this sexual reproduction event remain unclear. Haploid males occur sporadically in *O. biroi* [26,36]. Assuming that at least some of these males are fertile, this opens the possibility of mating and sexual reproduction. However, *O. biroi* lacks a queen caste, and *O. biroi* workers lack a spermatheca [25], implying that they are incapable of mating and storing sperm. A plausible explanation would be that at some point, clonal line A produced a mating-competent female (perhaps a queen), and that this female mated with a haploid male from clonal line B (**figure 2**). This would imply that *O. biroi* (or at least clonal line A) has the latent capacity to produce queens or other mating-competent females. However, such individuals have yet to be described.

**figure 2.**
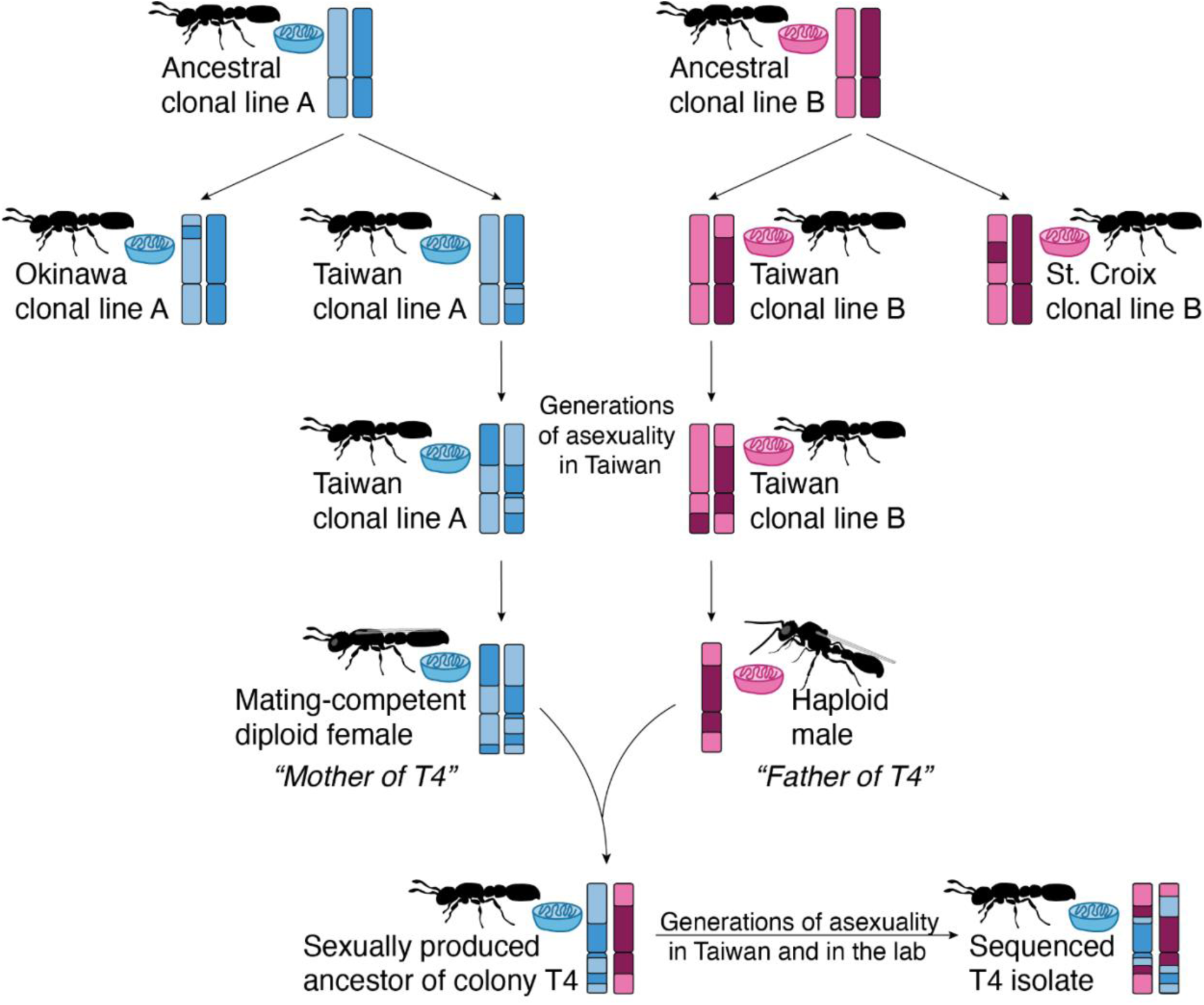
Model for the sexual origin of clonal line T4. Coloring on chromosomes and mitochondria indicates genetic identity, with blue for clonal line A and red for clonal line B. Dark and light shading on chromosomes indicates allelic identity.

Based on the co-occurrence of clonal lines A, B, and colony T4 in Taiwan, it is tempting to speculate that the sexual reproduction event that gave rise to the ancestor of T4 occurred in Taiwan after clonal lines A and B had been introduced there. This would imply that sexual reproduction can happen in the invasive range of *O. biroi*, whereas only asexual reproduction had been documented in the invasive range previously [26]. However, it is also possible that this sexual reproduction event occurred elsewhere in the invasive range, or even in the native range, with clonal line T4 being subsequently transported to Taiwan. Thus, we cannot firmly conclude that sexual reproduction occurs outside of the native range of *O. biroi*.

When did this sexual reproduction event occur? The colony was collected in 2001, and it would have taken at least a few years for the colony to grow to a noticeable size from the sexually produced founding individual. Therefore, at the latest, the event could have occurred in the late 1990s. Then, based on our mitochondrial mutation rate estimates, it may have occurred as early as the 1620s. Although we caution that this is a rough estimate that relies on many assumptions, this range encompasses current estimates of when *O. biroi* began to spread across the globe from its native Bangladesh via human commerce [33]. This implies that, despite *O. biroi* reproducing nearly entirely asexually, sex may have occurred sometime in the past few hundred years and may still occur sporadically today.

Rare sex implies that *O. biroi* might receive the benefits of both asexual and sexual reproduction. By creating new combinations of alleles, even rare sex can restore much of the lost adaptive potential to asexual lineages and allow for the purging of deleterious alleles [18,19,50,51]. In combination with high-fidelity maintenance of heterozygosity during asexual reproduction [30], rare sex could provide enough benefit to allow the long-term persistence of asexuality in *O. biroi*.

## Data accessibility

All DNA sequencing data are publicly available at the National Center for Biotechnology Information Sequence Read Archive under accession number PRJNA1277357. All code is available on GitHub (https://github.com/Social-Evolution-and-Behavior/TheOneWhereClonalAntsHadSex).

## Authors’ contributions

KDL: conceptualization, data curation, formal analysis, investigation, methodology, software, visualization, writing—original draft, writing—review & editing; NC: resources, writing—review & editing; DJCK: conceptualization, funding acquisition, project administration, resources, supervision, validation, writing—original draft, writing—review & editing.

## Acknowledgements

We thank Stephany Valdés-Rodríguez, Leonora Olivos-Cisneros, Alejandra Hurtado-Giraldo, and other members of the Kronauer Laboratory for ant maintenance and for valuable discussions; Connie Zhao and the Rockefeller University Genomics Resource Center for Illumina sequencing; Tanja Schwander, Tiphaine Bailly, and Yukina Chiba for comments on an earlier version of the manuscript. This is Clonal Raider Ant Project paper number 48.

---

**figure S1.**
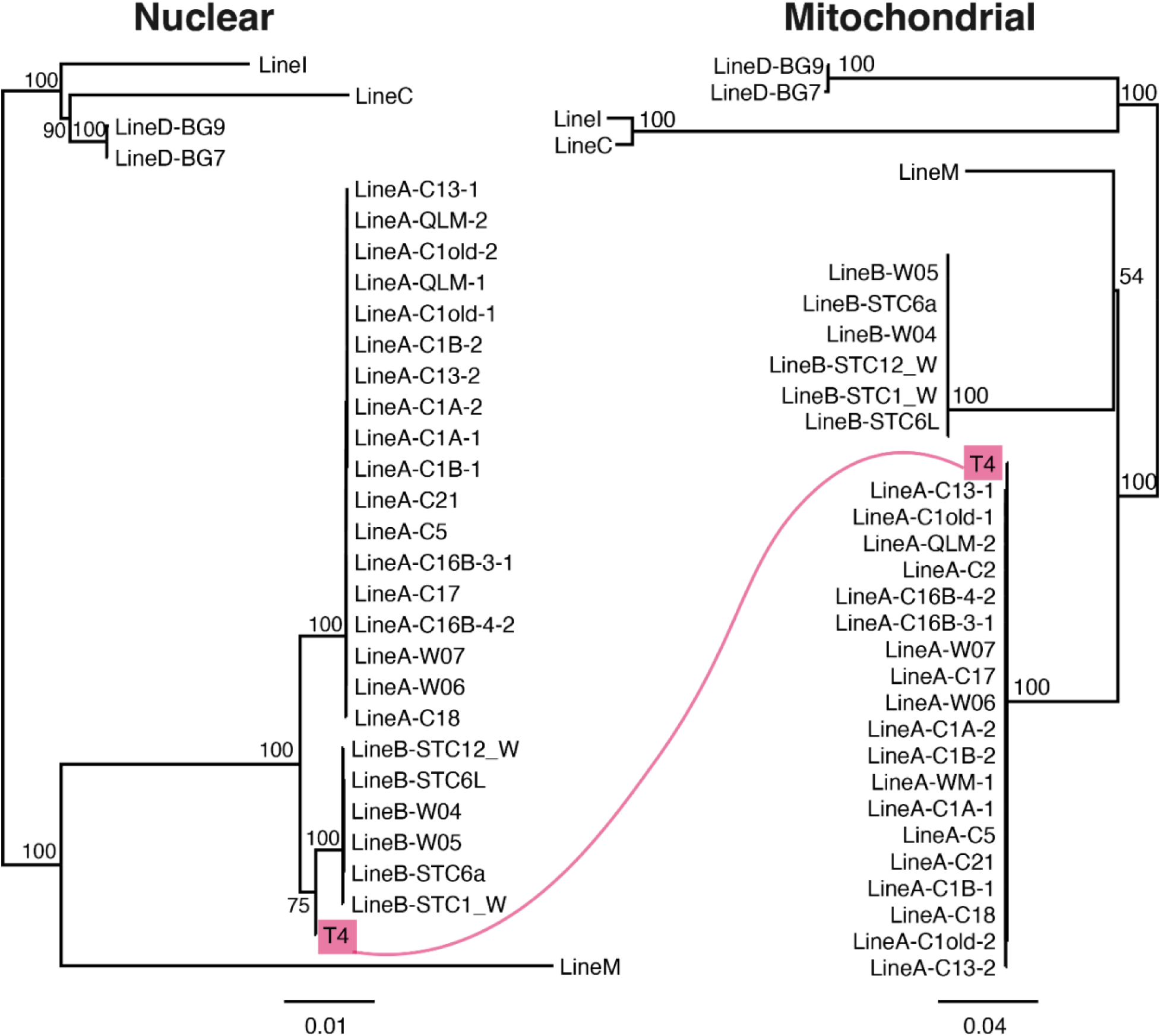
Nuclear and mitochondrial maximum likelihood phylogenies for all sequenced *O. biroi* samples. Midpoint-rooted maximum likelihood phylogenies based on SNPs from nuclear and mitochondrial genomes. Bootstrap supports from 1000 bootstrap replicates for major nodes are shown. Scale bars indicate substitutions per site.

**figure S2.**
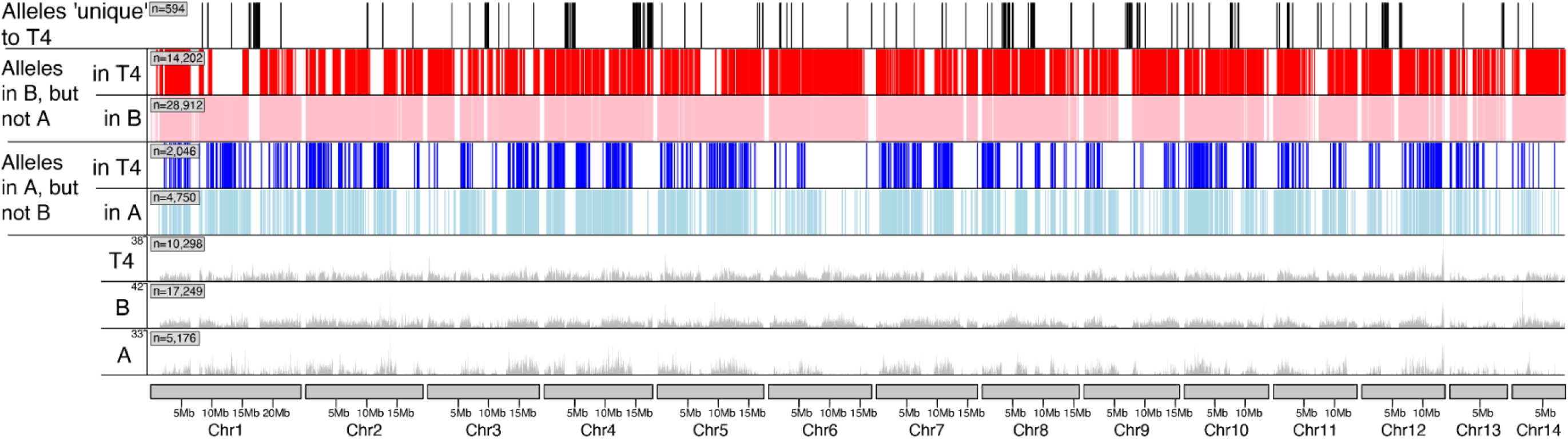
Private alleles from clonal lines A and B and their presence in colony T4. Karyoplot depicting, for all chromosomes in the *O. biroi* genome, sites with informative alleles in clonal line A, clonal line B, and colony T4, all shown as vertical tick marks. Heterozygous sites are shown in the gray histograms. For each plot, the number of variants (alleles) is shown in a gray box above chromosome 1.

**figure S3.**
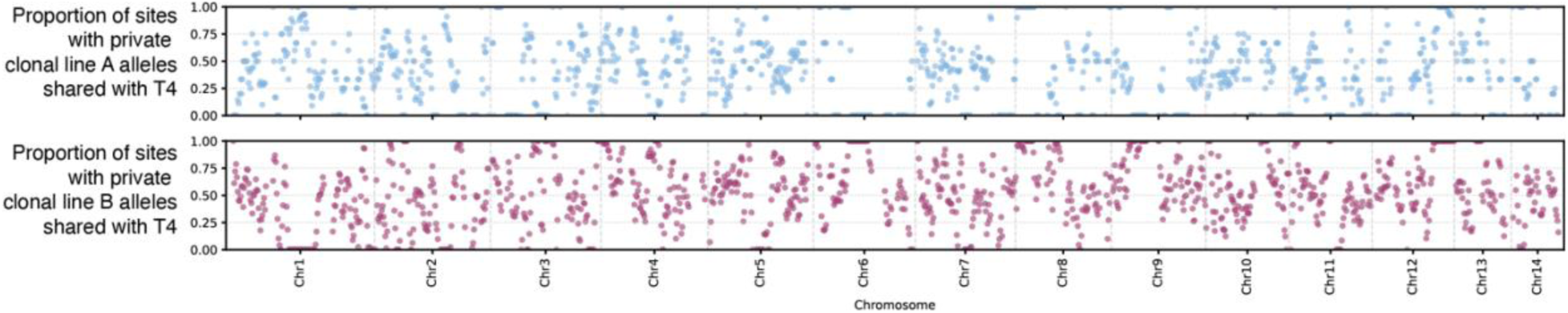
Proportions of private alleles from clonal lines A and B that are present in colony T4. Scatterplot depicting, for 600kb windows sliding every 150kb, the proportion of sites with alleles that are otherwise private to clonal line A or B that are shared with colony T4.

**table S1.** Metadata for DNA sequencing libraries used in this study.

| Sample Name | Clonal Line | Stock Colony | Sex | Ploidy | Published in Study | BioProject |
| --- | --- | --- | --- | --- | --- | --- |
| BG9_W | D | BG9 | Female | Diploid | Ref. 36 | PRJNA1075055 |
| C13-1 | A | C13 | Female | Diploid | Ref. 35 | PRJNA923657 |
| C13-2 | A | C13 | Female | Diploid | Ref. 35 | PRJNA923657 |
| C16B-3-1 | A | C16 | Female | Diploid | Ref. 30 | PRJNA947942 |
| C16B-4-2 | A | C16 | Female | Diploid | Ref. 30 | PRJNA947942 |
| C17 | A | C17 | Female | Diploid | Ref. 35 | PRJNA923657 |
| C18 | A | C18 | Female | Diploid | Ref. 35 | PRJNA923657 |
| C1A-1 | A | C1 | Female | Diploid | Ref. 35 | PRJNA923657 |
| C1A-2 | A | C1 | Female | Diploid | Ref. 35 | PRJNA923657 |
| C1B-1 | A | C1 | Female | Diploid | Ref. 35 | PRJNA923657 |
| C1B-2 | A | C1 | Female | Diploid | Ref. 35 | PRJNA923657 |
| C1old-1 | A | C1 | Female | Diploid | Ref. 35 | PRJNA923657 |
| C1old-2 | A | C1 | Female | Diploid | Ref. 35 | PRJNA923657 |
| C21 | A | C21 | Female | Diploid | Ref. 35 | PRJNA923657 |
| C5 | A | C5 | Female | Diploid | Ref. 35 | PRJNA923657 |
| W06 | A | C16 | Female | Diploid | Ref. 35 | PRJNA923657 |
| W07 | A | C16 | Female | Diploid | Ref. 35 | PRJNA923657 |
| W04 | B | STC6 | Female | Diploid | Ref. 30 | PRJNA947942 |
| W05 | B | STC6 | Female | Diploid | Ref. 30 | PRJNA947942 |
| LineC | C | Unknown | Female | Diploid | Ref. 36 | PRJNA1075055 |
| LineD | D | Unknown | Female | Diploid | Ref. 36 | PRJNA1075055 |
| LineI | I | Unknown | Female | Diploid | Ref. 36 | PRJNA1075055 |
| LineL | M | BG14 | Female | Diploid | Ref. 36 | PRJNA1075055 |
| MD-B-1-D | B | STC6 | Female | Diploid | Ref. 30 | PRJNA947942 |
| MD-B-1-M | B | STC6 | Female | Diploid | Ref. 30 | PRJNA947942 |
| MD-B-7-D1 | B | STC6 | Female | Diploid | Ref. 30 | PRJNA947942 |
| MD-B-7-M | B | STC6 | Female | Diploid | Ref. 30 | PRJNA947942 |
| STC1_W | B | STC1 | Female | Diploid | Ref. 30 | PRJNA947942 |
| STC12_W | B | STC12 | Female | Diploid | Ref. 30 | PRJNA947942 |
| STC6a | B | STC6 | Female | Diploid | Ref. 36 | PRJNA1075055 |
| STC6L | B | STC6 | Female | Diploid | Ref. 30 | PRJNA947942 |
| T4 | T4 | T4 | Female | Diploid | This study | PRJNA1277357 |
| WM-1 | A | QLM | Female | Diploid | Ref. 35 | PRJNA923657 |
| WM-2 | A | QLM | Female | Diploid | Ref. 35 | PRJNA923657 |

**table S2.** Cross-validation (CV) error estimates for ADMIXTURE runs across K values. CV error values for each run with different K values (numbers of assumed ancestral populations), with lower values suggesting better model fit. The minimum CV error was observed at K = 2, supporting the presence of two major genetic clusters in the dataset.

| **K** | **CV Error** |
| --- | --- |
| 1 | 0.40536 |
| 2 | 0.04607 |
| 3 | 0.07496 |
| 4 | 0.08966 |
| 5 | 0.16603 |
| 6 | 0.16092 |

